# Brain network interconnectivity dynamics explain metacognitive differences in listening behavior

**DOI:** 10.1101/2023.07.11.548535

**Authors:** Mohsen Alavash, Jonas Obleser

## Abstract

Complex auditory scenes pose a challenge to attentive listening, rendering listeners slower and more uncertain in their perceptual decisions. How can we explain such behaviors from the dynamics of cortical networks that pertain to the control of listening behavior? We here follow up on the hypothesis that human adaptive perception in challenging listening situations is supported by modular reconfiguration of auditory-control networks in a sample of N=40 participants (13 males) who underwent resting-state and task functional magnetic resonance imaging (fMRI). Individual titration of a spatial selective auditory attention task maintained an average accuracy of∼ 70% but yielded considerable inter-individual differences in listeners’ response speed and reported confidence in their own perceptual decisions. Whole-brain network modularity increased from rest to task by reconfiguring auditory, cinguloopercular, and dorsal attention networks. Specifically, interconnectivity between the auditory network and cinguloopercular network decreased during the task relative to the resting state. Additionally, interconnectivity between the dorsal attention network and cinguloopercular network increased. These interconnectivity dynamics were predictive of individual differences in response confidence, the degree of which was more pronounced after incorrect judgments. Our findings uncover the behavioral relevance of functional crosstalk between auditory and attentional-control networks during metacognitive assessment of one’s own perception in challenging listening situations and suggest two functionally dissociable cortical networked systems that shape the considerable metacognitive differences between individuals in adaptive listening behavior.

**Significance Statement:** The ability to communicate in challenging listening situations varies not only objectively between individuals but also in terms of their subjective perceptual confidence. Using fMRI and a challenging auditory task, we demonstrate that this variability in the metacognitive aspect of listening behavior is reflected on a cortical level through the modular reconfiguration of brain networks. Importantly, task-related modulation of interconnectivity between the cinguolopercular network and each auditory and dorsal attention network can explain for individuals’ differences in response confidence. This suggests two dissociable cortical networked systems that shape the individual evaluation of one’s own perception during listening, promising new opportunities to better understand and intervene in deficits of auditory perception such as age-related hearing loss or auditory hallucinations.

## Introduction

In complex auditory scenes, attending only to what we want to hear can be a challenging task, making our listening prone to errors and uncertainties. Listening as such not only hinges on auditory fidelity but also requires brain networks associated with directing, switching, and maintaining auditory attention (Shinn-Cunningham, 2008; Hill and Miller, 2010). In recent years there has been growing interest in studying listening as a trait-like behavior, as individuals differ substantially in utilizing cognitive strategies to control their auditory attention (Peelle, 2017; Shinn-Cunningham, 2017; Waschke et al., 2017; Tune et al., 2021). Nevertheless, the neural underpinnings of this inter­ individual variability are poorly understood.

Previous functional magnetic resonance imaging (fMRI) studies have associated auditory attention with regions overlapping with temporal and dorsal-parietal cortices (Puschmann et al., 2017; Shiell et al., 2018) and cinguloopercular/insular cortices in challenging listening tasks (Eckert et al., 2009; Wild et al., 2012; Erb et al., 2013; Vaden et al., 2019). Additionally, as reviewed elsewhere, when listeners selectively focus their auditory attention to a particular location, cortical areas overlapping with well-known visuospatial attention networks are more active compared to when not listening (Lee et al., 2014).

In parallel, functional connectivity studies have identified large-scale brain networks involved in cognitively demanding tasks, which include a cinguloopercular/insular network (Dosenbach et al., 2008; Menon and Uddin, 2010; Sadaghiani and D’Esposito, 2015), a frontoparietal network (Seeley et al., 2007; Marek and Dosenbach, 2018), and a dorsal attention network (Fox et al., 2006; Szczepanski et al., 2013). Surprisingly, whether and how these attentional control networks support adaptive listening behavior has remained somewhat understudied and incompletely understood.

The emerging field of network neuroscience provides the theoretical basis to address this question (Bassett and Sporns, 2017). Along these lines of research, our previous fMRI study provided a large-scale brain network account of adaptive listening behavior (Alavash et al., 2019). Specifically, we were able to explain individual adaptation to a challenging listening task with concurrent speech by reconfiguration of a network of auditory, cinguloopercular, and ventral attention nodes toward increased modular segregation during the task relative to resting-state baseline.

However, an often understudied and thus not explained aspect of human adaptive perception and communication lies in the subjective assessment of one’s own perceptual judgments. Many neuroimaging studies of auditory cognition, including our own, have measured behavior only objectively using individuals’ response accuracy (also labeled as “type I” perceptual decisions; Galvin et al., 2003) and, sometimes, speed. Challenging listening situations naturally introduce uncertainties to individual’s evaluation of their own auditory perception, thus confidence in listening (secondary to listening accuracy). This subjective evaluation can be viewed as a second-order measure of listening behavior, also referred to as “type II” decision (see above ref.). Such subjective reports can vary markedly from objective measures of listening behavior (Wostmann et al., 2021).

Thus, one important question is whether and how brain network dynamics regulate listener’s metacognitive assessment of their own perception to support adaptive listening. Knowing the role of cinguloopercular and insular networks in listening under challenging situations, performance monitoring, and interoception (Fleming and Dolan, 2012; Barttfeld et al., 2013; Uddin, 2015; Rouault et al., 2018), it is plausible to associate these interoceptive networks with subjective evaluation of one’s own perception during listening, that is, with a variant of “meta-listening”. Specifically, we do not know whether and how listeners’ capacity for introspection to reflect on their own perception relies on the interactions between distributed cortical networks.

To address this issue, we aimed to identify and characterize auditory-attentional control networks and their reconfiguration in a challenging auditory task that allowed joint measurement of objective and subjective listening behavior. Thus, leveraging the degree and direction of network dynamics, the present study asks two questions: (1) how are large-scale cortical networks being reconfigured to support selective listening? and (2) how can such reconfigurations explain individual differences not only in objective-perceptual, but also in subjective-metacognitive measures of listening behavior?

## Materials and Methods

### Participants

Forty-five participants were invited to take part in the study. All participants were right-handed (assessed by a translated version of the Edinburgh Handedness Inventory; Oldfield, 1971), had normal or corrected-to-normal vision, and all had normal hearing (assessed using pure tone audiometry (PTA); left-right average PTA across frequencies .025-8KHZ < 25 dbHL). None of the participants had any history of neurological or psychiatric disorders. Five participants were excluded as their data did not fulfill the criteria used to assess the quality of fMRI images (see section *FMRI data acquisition and preprocessing* below). Accordingly, 40 participants were included in the main analysis (age range = 18-32 years, median age = 23, 13 males). All procedures were in accordance with the Declaration of Helsinki and approved by the local ethics committee of the University of Lübeck. All participants gave written informed consent and were financially compensated (10€/h).

### Procedure

Each imaging session consisted of seven fMRI runs (Fig. 1A): eyes-open resting state (∼10 min) followed by six runs in which participants performed an adaptive selective pitch discrimination task (∼10 min each). During each resting state and each run of the listening task, 610 and 550 functional volumes were acquired, respectively. A structural scan (∼5 min) was acquired at the end of the imaging session. Before functional and structural imaging, outside the MRI room, each participant completed a screening procedure used to assess handedness, a shortened version of the Spatial, Speech and Hearing Quality Scale (SSQ; Gatehouse and Noble, 2004), as well as complete pure tone and speech-in-noise audiometry in free field in a sound-attenuated cabin.

**Figure 1.**
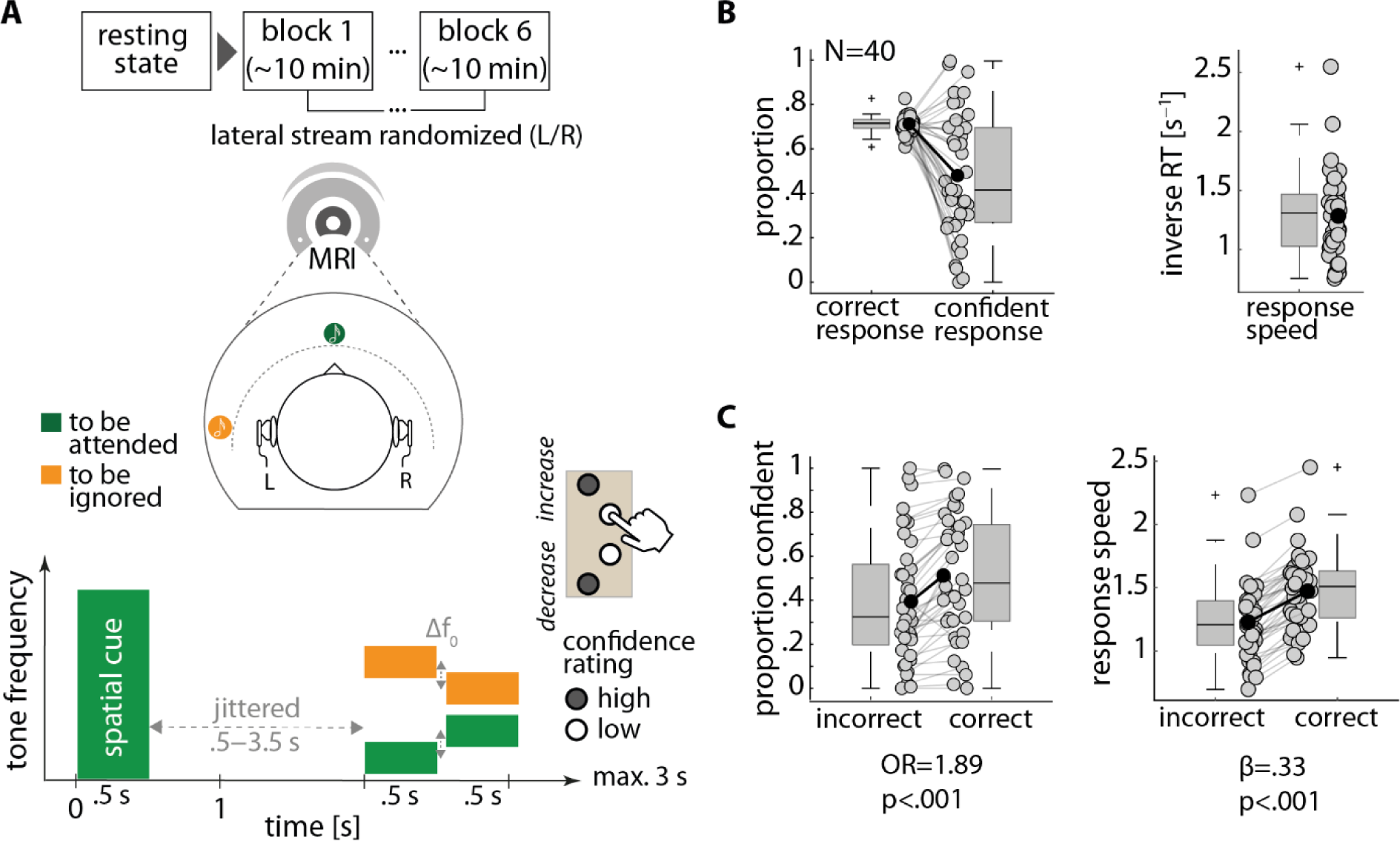
Listening paradigm and individuals’ behavior. **(A)** Functional MRI experiment. Following a 10-min eyes-open resting state, participants performed six blocks of purely auditory selective pitch discrimination task. During the task, two competing tone sequences were presented using in-ear headphones. The virtual spatialization of sounds was achieved using individually selected head-related transfer functions (HRTFs). One tone sequence was always presented at front and the other was switched between left and right over task blocks (left/right order randomized across participants). Each trial started by presentation of a broadband auditory cue {0-10 kHz; 0.5 s) followed by a jittered silent interval. Next, two tone-sequences, each consisting of two brief (0.5 s) complex tones, were presented at different locations (front, left, or right). Fundamental frequencies of low-frequency tones within each sequence were fixed at 177 and 267 Hz. Frequencies of high­ frequency tones were titrated throughout the experiment (Mo: tracking parameter). Participants had to judge whether the target tone sequence at the cued location had increased or decreased in pitch. Participants also reported their confidence in their own decision (high: outer buttons; low: inner buttons). **(B)** Individuals’ listening behavior. Listeners’ behavior was measured both objectively and subjectively using response accuracy, speed (i.e., inverse response time; 1/RT), and confidence rating. Individuals showed considerable variability in their listening behavior, notably in response speed and confidence. **(C)** Prediction of single-trial response confidence and speed from accuracy. Listeners were more confident and faster in their responses on correct trials as compared to incorrect trials (linear mixed-effects models with single-trial accuracy as a binary predictor). *Box plots.* The central line indicates the median, and the bottom and top edges of the box indicate the 25th and 75th percentiles, respectively. The whiskers extend to the most extreme data points not considered outliers. The outliers are marked by’+’ symbol. *Individual data.* The central solid circle indicates the mean.

### Experimental design

Task design and stimuli were adapted from Dai et al. (2018) and were identical to our previous EEG study (Wöistmann et al., 2019). The experiment was implemented in the Psychtoolbox for Matlab (Brainard, 1997), and conducted inside the MRI scanner with the participant lying in the supine position (Fig. 1A).

#### Auditory stimuli

All auditory stimuli were presented at a sampling frequency of 44.1 kHz at a comfortable level of ∼65 dBA. On each trial, an auditory spatial cue (10.9 kHz low-pass filtered Gaussian noise; 0.5 s) was presented at one location followed by two concurrent tone sequences presented at two different spatial locations (front, left, or right). Each tone sequence consisted of two 0.5 s complex tones (fixed ISi of 50 ms), one low-pitch tone and one high-pitch tone. All tones and the spatial cue were gated on and off with 100 ms cosine ramps. The pitch of the low-pitch tone was fixed at 177 Hz (including 32 harmonics) for one sequence and at 267 Hz (including 2 harmonics) for the other sequence. Throughout the experiment, the pitch difference was individually titrated by varying the fundamental frequency of the high-pitch tone in each sequence in semitones relative to the low-pitch tone using an adaptive tracking procedure (two up, one down) to retain the listener’s overall accuracy at∼ 70% (Levitt, 1971; see Fleming et al., 2010, and Palmer et al., 2014, for a similar titration approach in a visual perceptual task). Thus, after one incorrect response or two subsequent correct responses the pitch difference within each tone sequence was increased or decreased in steps of 0.05 semitones on the next trial, respectively. This procedure ensured that the task was equally challenging enough, as reflected in a fair number of trials with correct and incorrect responses for each listener, and in turn yielded considerable inter-individual variability in response speed and confidence across listeners (Fig. 1B).

The initial pitch difference for the tracking procedure was obtained from a pre-experiment training session. The cue location (front vs. side), the pitch direction within each sequence (increasing vs. decreasing), and the assignment of tone sequences to the locations were balanced across trials and drawn randomly for an individual trial. Tone sequences were presented using in-ear headphones based on individually selected head-related transfer functions. The location of a sound stream could be either front or side (i.e., 0 or 90° azimuth relative to ear-nose-ear line). The location of the lateral stream changed between left and right across blocks of the experiment. The other stream was always presented at the front throughout. There were three blocks of the experiment with the lateral stream on the left side, and three blocks with the lateral stream on the right. The order of blocks was counterbalanced across participants and alternated between blocks with the lateral stream on the left versus right. Within each block, a participant completed 96 trials (each spatial location served as the target in 48 trials).

#### Virtual spatialization of sounds

To present the auditory stimuli used in the task (i.e., the spatial cue and complex tones) in three different spatial locations (i.e., left, right, front), the stimuli were convolved in Matlab during the presentation in the Psychtoolbox environment with individually selected head-related transfer functions (HRTFs) from the publicly available HRTF database CIPIC (https://github.com/amini-allight/cipic-hrtf-database). A similar procedure has been used in previous studies (e.g., Shiel et al., 2018). The procedure to select individual HRTF was conducted before training, outside the MRI scanner room, and using the same in-ear headphones as the main experiment. This procedure was completed in two steps. First, in a pre-selection step, individual anthropometric data were measured and used to find the HRTF which would match the best to each individual based on the Euclidean distance as the similarity measure. Next, these best matches were sorted and used to present the spatial cue at the front without elevation three times. Finally, the participant was asked whether the stimulus gave the impression that the cue was presented at the front, in the middle and outside the head. If not, the second step was repeated using the next impulse response in order, until the participant confirmed the virtual specialization of the cue at the front.

#### Selective pitch discrimination task

At the start of each trial, after a jittered period of ∼1 s (0.8-1.2 s), an auditory spatial cue was presented on one location to inform the participant about the target location (Fig. 1A). After a jittered period of ∼1.5 s (right-skewed distribution; median: 1.44 s; truncated at .5 and 3.5 s) relative to cue offset, two tone sequences were presented concurrently. Participants reported whether the tone sequence at the target location increased or decreased in pitch and how confident they were in this response using a response box with four buttons. No visual stimulation was done during the experiment, and the participants were instructed to fixate on a cross in the middle of a screen with a gray background projected from the back of the magnet bore onto a mirror mounted on the head coil and positioned in front of the participant’s view. Before the main experiment, a short training ensured that participants could perform the pitch discrimination task.

### FMRI data acquisition and preprocessing

#### MRI image acquisition parameters

Functional MRI data were collected using a Siemens MAGNETOM Skyra 3T scanner equipped with a 64-channel head/neck coil and an echo-planar imaging (EPI) sequence [repetition time (TR) =1000 ms; echo time (TE) =25 ms; flip angle (FA) =60°; acquisition matrix =64x64; field of view (FOV) =192 mm x 192 mm; voxel size =3x3x3 mm; slice spacing =3 mm]. Each image volume consisted of 36 oblique axial slices parallel to the anterior commissure-posterior commissure (AC-PC) line and was acquired with an acceleration factor of 2. Structural images were obtained using a magnetization prepared rapid gradient echo (MP-RAGE) sequence [TR =1900 ms; TE =2.44 ms; FA =9°; 1-mm isotropic voxel; 192 sagittal slices].

#### MRI image preprocessing

An overview of the preprocessing steps for analyzing the fMRI data is presented in Fig. 2. Volumetric images acquired from each participant were converted from DICOM format to standard BIDS format (Gorgolewski et al., 2016) using HeuDiConv (Halchenko, 2018). The resulting data were then preprocessed using fMRIPrep (Esteban et al., 2019) with minimal preprocessing applied according to the standard fMRIPrep procedure. Results included in this manuscript come from preprocessing performed using *fMR/Prep* 20.2.6 (Esteban, Markiewicz, et al. 2018; Esteban, Blair, et al. 2018), which is based on *Nipype* 1.7.0 (Gorgolewski et al. 2011; Gorgolewski et al. 2018).

**Figure 2.**
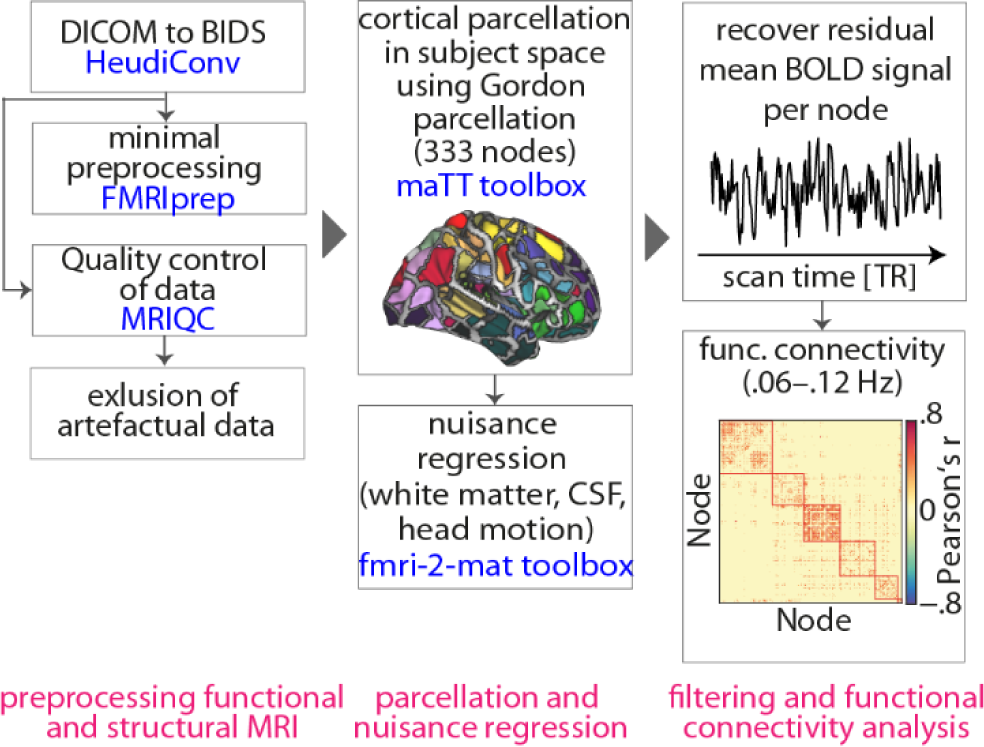
Analysis steps through which artefact-clean mean regional BOLD signals were recovered per cortical parcel. The results were used to construct graph-theoretical model of the whole-brain network during resting state and the listening task. Each step required publicly available toolboxes which are indicated in blue text.

#### Anatomical data preprocessing

The Tl-weighted (T1w) image was corrected for intensity non­ uniformity (INU) with N4BiasFieldCorrection (Tustison et al. 2010), distributed with ANTs 2.3.3 (Avants et al. 2008), and used as T1w-reference throughout the workflow. The T1w-reference was then skull-stripped with a *Nipype* implementation of the antsBrainExtraction.sh workflow (from ANTs), using OASIS3OANTs as target template. Brain tissue segmentation of cerebrospinal fluid (CSF), white-matter (WM) and gray-matter (GM) was performed on the brain-extracted T1w using fast (FSL 5.0.9, Zhang et al. 2001). Brain surfaces were reconstructed using recon-all (FreeSurfer 6.0.1, Dale, Fischl, and Sereno 1999), and the brain mask estimated previously was refined with a custom variation of the method to reconcile ANTs-derived and FreeSurfer-derived segmentations of the cortical gray-matter of Mindboggle (Klein et al. 2017). Volume-based spatial normalization to one standard space (MNl152NLin2009cAsym) was performed through nonlinear registration with ants Registration (ANTs 2.3.3), using brain-extracted versions of both Tl w reference and the Tl w template. The following template was selected for spatial normalization: *ICBM 152 Nonlinear Asymmetrical template version 2009c* (Fonov et al. 2009, TemplateFlow ID: MNI152NLin2009cAsym).

#### Functional data preprocessing

First, a reference volume and its skull-stripped version were generated using a custom methodology of *fMRIPrep.* Susceptibility distortion correction (SOC) was omitted. The BOLD reference was then co-registered to the Tl w reference using bbregister (FreeSurfer) which implements boundary-based registration (Greve and Fischl 2009). Co-registration was configured with six degrees of freedom. Head-motion parameters with respect to the BOLD reference (transformation matrices, and six corresponding rotation and translation parameters) are estimated before any spatiotemporal filtering using mcflirt (FSL 5.0.9, Jenkinson et al. 2002). BOLD runs were slice-time corrected to 0.462s (0.5 of slice acquisition range 0s-0.925s) using 3dTshift from AFNI 20160207 (Cox and Hyde 1997, RRID:SCR_005927). The BOLD time-series (including slice-timing correction when applied) were resampled onto their original, native space by applying the transforms to correct for head-motion. These resampled BOLD time-series will be referred to as *preprocessed BOLD in original space,* or just *preprocessed BOLD.* Several confounding time-series were calculated based on the *preprocessed BOLD:* framewise displacement (FD), DVARS and three region­ wise global signals. FD was computed using two formulations following Power (absolute sum of relative motions, Power et al. 2014) and Jenkinson (relative root mean square displacement between affines, Jenkinson et al. 2002). FD and DVARS are calculated for each functional run, both using their implementations in *Nipype* (following the definitions by Power et al. 2014). The three global signals are extracted within the CSF, the WM, and the whole-brain masks. Additionally, a set of physiological regressors were extracted to allow for component-based noise correction *(CompCor,* Behzadi et al. 2007). Principal components are estimated after high-pass filtering the *preprocessed BOLD* time­ series (using a discrete cosine filter with 128s cut-off) for the two *CompCor* variants: temporal (tCompCor) and anatomical (aCompCor). tCompCor components are then calculated from the top 2% variable voxels within the brain mask. For aCompCor, three probabilistic masks (CSF, WM, and combined CSF+WM) are generated in anatomical space. The implementation differs from that of Behzadi et al. in that instead of eroding the masks by 2 pixels on BOLD space, the aCompCor masks are subtracted a mask of pixels that likely contain a volume fraction of GM. This mask is obtained by dilating a GM mask extracted from the FreeSurfer’s *aseg* segmentation, and it ensures components are not extracted from voxels containing a minimal fraction of GM. Finally, these masks are resampled into BOLD space and binarized by thresholding at 0.99 (as in the original implementation). Components are also calculated separately within the WM and CSF masks. For each CompCor decomposition, the *k* components with the largest singular values are retained, such that the retained components’ time series are sufficient to explain 50 percent of variance across the nuisance mask (CSF, WM, combined, or temporal). The remaining components are dropped from consideration. The head-motion estimates calculated in the correction step were also placed within the corresponding confounds file. The confound time series derived from head motion estimates and global signals were expanded with the inclusion of temporal derivatives and quadratic terms for each (Satterthwaite et al. 2013). Frames that exceeded a threshold of 0.5 mm FD or 1.5 standardised DVARS were annotated as motion outliers. All resamplings can be performed with *a single interpolation step* by composing all the pertinent transformations (i.e., head-motion transform matrices, susceptibility distortion correction when available, and co-registrations to anatomical and output spaces). Gridded (volumetric) resamplings were performed using antsApplyTransforms (ANTs), configured with Lanczos interpolation to minimize the smoothing effects of other kernels (Lanczos 1964). Non-gridded (surface) resamplings were performed using mri_vol2surf (FreeSurfer). Many internal operations of *fMR/Prep* use *Ni/earn* 0.6.2 (Abraham et al. 2014), mostly within the functional processing workflow.

#### Data quality assessment

Image quality measures were obtained from the outputs of DICOM­ to-BIDS conversion using the MRIQC tool (Esteban et al., 2017). Data quality was assessed following the criteria outlined by Faskowitz et al. (2020). Specifically, functional data were excluded if more than 25% of the frames exceeded 0.2-mm framewise displacement (Parkes et al., 2018) or if marked as an outlier (exceeding 1.Sxinterquartile range (IQR) in the adverse direction) in more than half of the following seven image quality metrics (calculated within datasets, across all functional acquisitions): dvars, tsnr, fd mean, aor, aqi, snr, and efc (refer to MRIQC documentation for definitions). Consequently, five datasets and two runs from two participants were excluded from the main analysis.

### Functional connectivity and network analysis

#### Cortical parcellation

Consistent with our prior study (Alavash et al., 2019), cortical nodes were delineated using a previously established functional parcellation (Gordon et al., 2016), known for its relatively higher accuracy compared to other parcellations (Arslan et al., 2017). This parcellation, comprising 333 cortical areas, is based on surface-based resting-state functional connectivity boundary maps. To obtain the Gordon333 parcellation in subject space, we utilized the maTT toolbox as described by Faskowitz et al. (2020). In brief, a Gaussian classifier surface atlas was employed for each participant used in conjunction with FreeSurfer’s mris_ca_label function to transfer the group average atlas to subject space based on individual surface curvature and sulcal patterns, resulting in a T1w space volume for each participant. For compatibility with functional data, the parcellation was resampled to 2-mm T1w space.

#### Nuisance regression

To mitigate the effects of spurious temporal correlations induced by physiological and movement artifacts, a nuisance regression procedure was implemented using the fmri-2-mqt toolbox as described by Faskowitz et al. (2020). In this procedure, each preprocessed BOLD image underwent linearly detrending, confound regression, and standardization. The confound regression method mirrored that of our prior study (Alavash et al., 2019) and included six motion estimates and their first-order derivatives, time series of the mean cerebrospinal fluid, and mean white matter signal. Notably, global signal regression was omitted from the regional time series, as the impact of global signal regression remains uncertain in the field (Murphy and Fox, 2016; Power et al., 2017). Importantly, our primary findings highlight the reconfiguration of an auditory attentional-control network, distinct from the predominantly sensorimotor disruption observed in the global signal map (Power et al., 2017). Following preprocessing and nuisance regression, residual mean BOLD time series were obtained for each node, performed for both resting-state and listening­ task data. To ensure signal equilibration, the first 10 volumes of resting state and each run of the listening task were discarded.

#### Functional connectivity

Residual time series underwent band-pass filtering using the maximum overlap discrete wavelet transform (Daubechies wavelet of length eight as implemented in the waveslim R package), focusing on the range of 0.06-0.12 Hz (wavelet scale three), consistent with our prior work (Alavash et al., 2019). Prior research has demonstrated that behavioral correlates of the functional connectome are best observed by analyzing low-frequency large-scale brain networks (Zhang et al., 2016). The choice of wavelet scale three aligns with previous findings indicating that cognitive task-related behavior predominantly correlates with changes in functional connectivity within this frequency range (Alavash et al., 2015b; Alavash et al., 2018). Pearson correlations between wavelet coefficients were computed to establish the association between each pair of cortical regions, resulting in one 333 x 333 correlation matrix per participant for each resting state and run of the listening task.

#### Network analysis

Brain graphs were constructed from the functional connectivity matrices by retaining the top 10% of connections according to the rank of their correlation strengths (van Wijk et al., 2010; van den Heuvel et al., 2017). This process yielded sparse binary undirected brain graphs at a fixed network density of 10, ensuring uniformity in density across participants, resting state, and listening task. Mean functional connectivity was computed as the average of the upper diagonal elements of the sparse connectivity matrix for each participant. Additionally, network modularity was estimated to characterize the configuration of large-scale brain networks at a macroscopic level (Rubinov and Sporns, 2010).

#### Network modularity

Modularity describes the decomposability of a network into non­ overlapping sub-networks, characterized by having relatively dense intra-connections and relatively sparse inter-connections. Rather than an exact computation, modularity of a given network is estimated using optimization algorithms (Lancichinetti and Fortunato, 2009; Steinhaeuser and Chawla, 2010). The extent to which a network partition exhibits a modular organization is measured by a quality function, the so-called *modularity index* (*Q*). We used a common modularity index originally proposed by (Newman, 2006), and employed its implementation in the Brain Connectivity Toolbox (Rubinov and Sporns, 2010) which is based on the modularity maximization algorithm known as Louvain (Blondel et al., 2008). The modularity index is defined as:

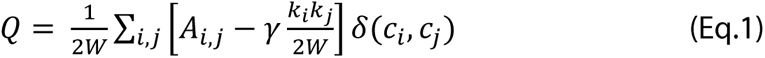

*Q* ranges between -1 and 1. In Eq. 1, *A_i,j_* represents the weight (zero or one if binary) of the links between node *i* and *j*, *k_i_* = ∑*_j_ A_i,j_* is the sum of the weights of the links connected to node *i*, and *c_i_* is the community or module to which node *i* belongs. The δ-function δ(*u v*) is 1 if *u* = *v* and 0 otherwise, and 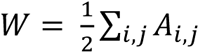. Similar to previous work (Bassett et al., 2010; Alavash et al., 2018), the structural resolution parameter γ (see Fortunato and Barthelemy, 2007; Lohse et al., 2014) was set to unity for simplicity. The maximization of the modularity index *Q* gives a partition of the network into modules such that the total connection weight within modules is as large as possible, relative to a commonly used null model whose total within-module connection weights follows 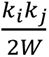. Thus, a “good” partition with *Q* closer to unity gives network modules with many connections within and only few connections between them; in contrast, a “bad” partition with *Q* closer to zero gives network modules with no more intra-module connections than expected at random (Good et al., 2010). Thus, higher *Q* reflects higher functional segregation on the intermediate level of network topology (Rubinov and Sporns, 2010; Betzel and Bassett, 2016). Due to stochastic initialization of the greedy optimization, the module detection algorithm was applied 100 times for each brain graph, and the highest *Q* value obtained was used as the modularity index in the subsequent statistical analyses.

#### Consensus modularity

Repetition of the module detection algorithm leads to multiple possible high-modularity partitions that maximize *Q* for a given network, resulting in module membership assignments that vary across runs of the algorithm (Good et al., 2010). In order to account for this variability, we used the consensus approach proposed by Lancichinetti and Fortunato (2012) whereby an agreement matrix is calculated, representing the probability of each node pair to be assigned to the same module across iterations. Finally, the agreement matrix was subjected to an independent module detection, resulting in an individual-level (or group-representative when the input was group-average connectivity matrix) high-modularity partition. In this step, the resolution parameter τ was set to 0.75, representing the level at which the agreement matrix was thresholded before being subjected to the final module detection. The modularity detection was implemented with no prior community affiliation input, hence in a purely data-driven fashion.

#### Group-level modularity partition

Based on the results obtained from graph-theoretical consensus community detection (see above), group-level functional connectivity per resting state and listening task was computed by first averaging unthresholded (raw) connectivity matrices across all participants, and then including the top 10% of the connections in the graph according to the rank of their correlation strengths (Ginestet et al., 2011; Fornito et al., 2013). To obtain group-level modularity partition, the graph-theoretical consensus community detection algorithm was applied to the sparse group-level connectivity matrix (identical to Bertolero et al., 2015). Similar to Gordon et al. (2016), modules with fewer than five nodes were removed, and the result was used for visualization. To functionally identify the network modules, we used the labels assigned to each cortical node in Gordon et al. (2016).

#### Network visualization

Brain surfaces were visualized using Connectome Workbench. Connectograms were visualized using Brain Data Viewer. Network flow diagrams were visualized using MapEquation.

### Statistical analysis

#### Behavioral data

The performance of participants in the listening task was evaluated based on three measures: correct pitch discrimination (response accuracy: correct/incorrect), response speed (the inverse of response time), and confidence rating (response confidence: high/low). Trials without a response within the response time window and those with response times less than 300 ms were excluded. Single-trial behavioral measures for each participant were considered as the dependent variable in the (generalized) linear mixed effects analysis. In models predicting confidence or speed, response accuracy was treated as a binary predictor.

#### Brain data

Statistical comparisons of connectivity and network modularity between resting state and the listening task were conducted using permutation tests for paired samples (the rest and task labels were randomly permuted 10,000 times). Cohen’s d for paired samples was used as the corresponding effect size.

#### Brain-behavior models

The relationship between brain activity and behavior was explored using linear mixed effects analysis. Each single-trial behavioral measure (accuracy, speed, or confidence) across participants was the dependent variable. In models predicting confidence or speed, response accuracy was used as a binary predictor. The main predictors included titrated pitch difference, time-on-task (task block number: 1-6), congruency of tone sequences (congruent vs. not congruent), location of the lateral stream (left vs. right), and the role of the lateral stream (attended vs. ignored). Give the study’s focus on the reconfiguration of resting-state brain networks from rest to task and its relation to listening behavior, mean connectivity or modularity of brain networks were used as main predictors. Specifically, the main effect of each rest and task connectivity (or modularity) was included in the models as between-subject covariates. Additionally, to account for activity levels in auditory, cinguloopercular, and dorsal attention network, mean beta estimates obtained from activation analysis were averaged within each network and included in the models as between-subject covariates.

#### Brain-behavior models

The relationship between brain activity and behavior was explored using linear mixed effects analysis. Each single-trial behavioral measure (accuracy, speed, or confidence) across participants was the dependent variable. In models predicting confidence or speed, response accuracy was used as a binary predictor. The main predictors included titrated pitch difference, time-on-task (task block number: 1-6), congruency of tone sequences (congruent vs. not congruent), location of the lateral stream (left vs. right), and the role of the lateral stream (attended vs. ignored). Give the study’s focus on the reconfiguration of resting-state brain networks from rest to task and its relation to listening behavior, mean connectivity or modularity of brain networks were used as main predictors. Specifically, the main effect of each rest and task connectivity (or modularity) was included in the models as between-subject covariates. Additionally, to account for activity levels in auditory, cinguloopercular, and dorsal attention network, mean beta estimates obtained from activation analysis were averaged within each network and included in the models as between-subject covariates.

Categorical predictors were coded using deviation coding, and continuous variables were Z­ scored. Generalized linear mixed effects models with the logit as a link function were used for accuracy and confidence, while linear mixed effects models with Gaussian distribution were used for speed. P values for individual model terms were derived using the Satterthwaite approximation for degrees of freedom and adjusted for the false discovery rate. Odds ratios (OR) were reported for accuracy or confidence models, and regression coefficients (β) for speed models. Analyses were conducted in R using the packages lme4 and sjPlot.

#### Bayes factor

To aid interpretation of significant and non-significant effects, Bayes Factors (BF) were calculated based on the comparison of Bayesian information criterion (BIC) values as proposed by Wagenmakers (2007). BF was calculated by comparing the BIC values of the full model to a reduced model without the accuracy ∈ interconnectivity interaction term: BF = exp([BIC(H0)­BIC(H1)]/2). Log-BFs larger than 1 provide evidence for the presence of an effect (i.e., the observed data are more likely under the more complex model) whereas log-BFs smaller than -1 provide evidence against the effect, following conventions outlined by Dienes (2014).

## Results

The auditory attention paradigm allowed listeners to judge the direction of pitch change in the to­ be-attended auditory object clearly above chance (with an accuracy of ∼70%) but left them uncertain about their own selective perceptual decisions, as reflected in an overall response confidence of around ∼50% (see Fig. 1B, left). Across listeners, the subjective assessment of one’s own perceptual decision varied considerably, as did listeners’ response speed (Fig. 1B, right).

Guided by our previous fMRI study on the brain network account of spatial selective auditory attention (Alavash et al., 2019), we predicted that during the task brain networks would reconfigure toward higher functional segregation as quantified by network modularity. Importantly, we asked whether such reconfiguration could explain inter-individual variability in listening behavior. To this end, we used (generalized) linear mixed effects models (GLMMs) to examine the influence of the experimental conditions and brain network dynamics on listeners’ single-trial behavior measured using response accuracy, speed, and confidence.

### High inter-individual variability in response speed and confidence during selective pitch discrimination

The average pitch difference between consecutive tones (i.e., the stimulus parameter tracked per listener adaptively; see Fig. 1A, *Mo)* was 0.84 semitones (± 0.69 semitones between-subject SD). As intended, the resulting accuracy (average proportion correct) was 71% (± 3.6% between-subject SD).

Notably, as illustrated in Fig.1B, listeners showed considerable inter-individual variability in their response speed (mean ± SD response speed = 1.23 ± 0.37 s-^1^; coefficient of variation = 0.3) and in their confidence (average proportion and SD of ‘confident’ responses = 48% ± 28%; coefficient of variation = 0.58).

Lending plausibility to the behavioral results obtained, single-trial listening behavior confirmed the beneficial effect of pitch difference on listeners’ behavior, i.e., the larger the pitch difference, the better the behavioral performance (GLMMs; accuracy: odds ratio (OR) = 1.42, p < 0.001; speed: = 0.26, p < 0.01; confidence: OR= 2.97, p < 0.001). Listeners also showed more accurate and faster performance when the direction of the pitch change was congruent across the two tone-sequences (accuracy: OR= 3.06, p < 0.001; speed: = 0.08, p < 0.01). In addition, listeners were less confident when the to-be-attended tone-sequence was presented in front compared with the side (OR= 0.84, p < 0.01). All of these effects were very well in line with the results we found in our previous EEG study in which the stimuli during the same task were presented in free field using loudspeakers (Wöistmann et al., 2019).

Of relevance to all further brain-behavior results, the three behavioral measures proved sufficiently reliable. Only if they are sufficiently reliable, meaning that individual data on repeated tests correlate positively, can any brain-behavior relation, which is the main scope here, be interpreted in a meaningful way (Guggenmos, 2021). Using the individual data across six blocks of the task, we calculated the reliability metric Cronbach’s alpha (CA) for each measure. The results indicated relatively high reliability for each behavioral measure, particularly for response speed and confidence (accuracy: CA= 0.58, speed: CA= 0.95, confidence: CA= 0.98).

Adaptive tracking of individual accuracy at∼ 70% throughout the experiment allowed us to split listeners’ single-trial responses into correct and incorrect judgments. Specifically, on average, listeners performed 486.89 ± 69.04 trials over six blocks of the task (excluding timeouts and trials with reaction times (RT) < 0.3 s or RT > 3 s relative to the onset of the second tone). Out of these, 346.19 ± 55.89 trials were correct and 133.6 ± 32.35 were incorrect. Accordingly, to investigate the relation between response accuracy and speed or confidence across listeners, we used single-trial accuracy as a binary predictor in linear mixed-effects models predicting single-trial response speed or confidence, separately. This analysis revealed that listeners were faster and more confident in their responses on correct trials as compared to incorrect trials (speed: = 0.33, p < 0.001; confidence: OR = 1.89, p < 0.001; see Fig. 1C). Important to the questions asked here, the individual ability to reflect on their own auditory perceptual decision varied markedly between listeners: while some listeners showed “overconfident” behavior (i.e., a high proportion of confident responses on incorrect trials), some showed “under-confident” behavior (i.e., a low proportion of confident responses on correct trials).

### Higher modularity of cortical networks during selective pitch discrimination

Using fMRI data, we initially investigated whether there is an increase in functional segregation of large-scale brain networks during the listening task. Our primary analysis involved comparing the functional segregation of brain networks during the task with that during rest. Functional segregation was assessed using modularity, a graph-theoretical measure capturing network decomposability into subsystems (Rubinov and Sporns, 2010; Fig. 3A). Modularity has been linked to cognitive flexibility and adaptive behaviors (Bassett et al., 2010; Alavash et al., 2015a; Bertolero et al., 2018). Building on our prior fMRI study (Alavash et al., 2019), we hypothesized an increase in brain network modularity during the selective pitch discrimination task compared to resting state.

**Figure 3.**
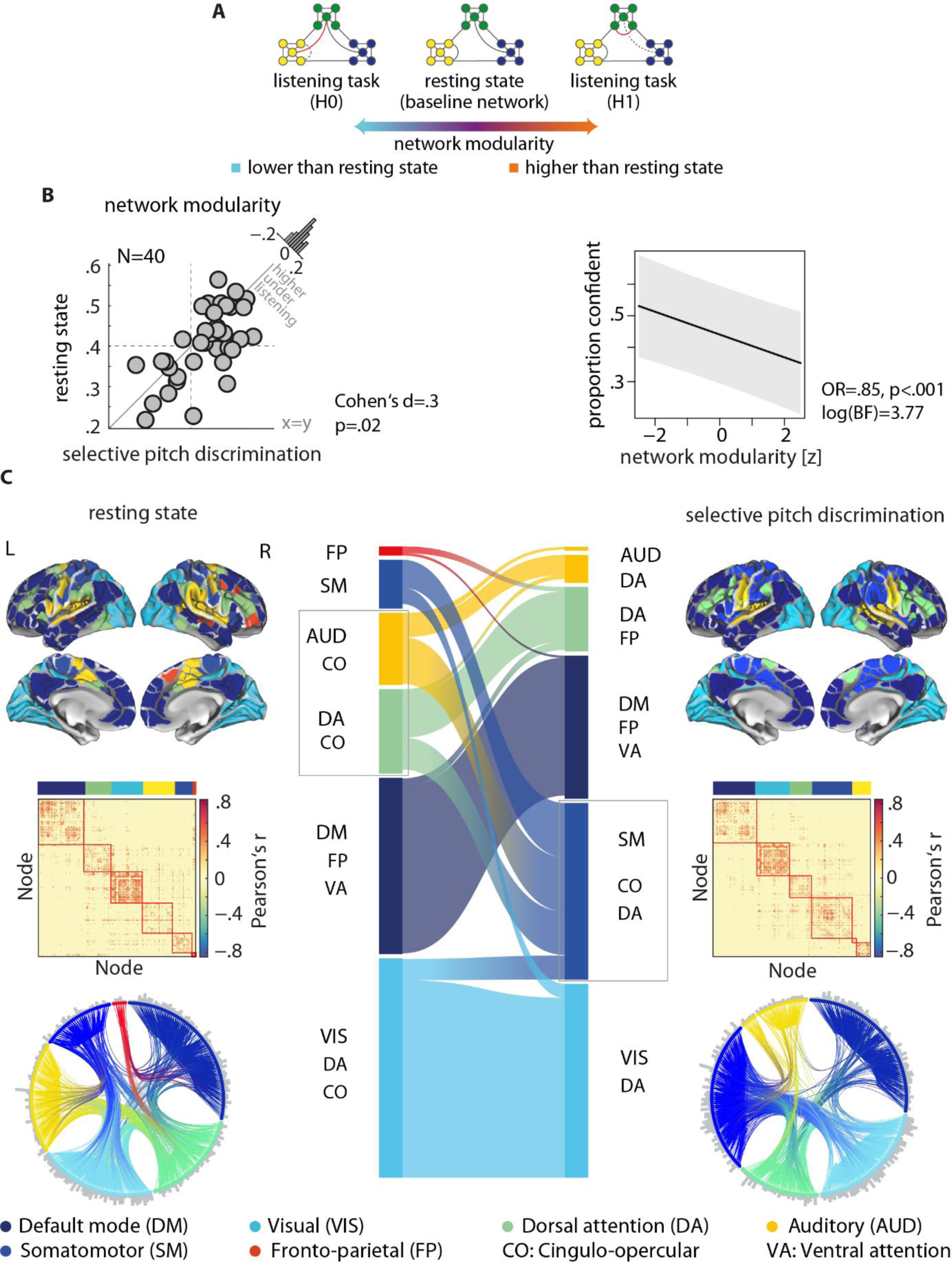
Reconfiguration of whole-brain network under the listening task relative to resting state. **(A)** Two possible network reconfigurations (i.e., left/right toy graphs compared to middle graph). Higher functional segregation is characterized by an increase in network modularity, a measure that quantifies grouping of nodes into relatively dense subsystems (here shown in distinct colors), which are sparsely interconnected. Toward higher functional segregation, the hypothetical baseline network loses the shortcut between the blue and green module (dashed link). Instead, within the green module, a new connection emerges (red link). (B) During the listening task, network modularity significantly increased (left plot), the degree of which predicted lower proportion of confident responses across listeners (right plot). Histograms show the distribution of the change (task minus rest) across all 40 listeners. (C) Whole-brain resting-state network decomposed into distinct modules shown in different colors. The network modules are visualized on the cortical surface, within the functional connectivity matrix, and on the connectogram using a consistent color scheme. Modules are identified according to their node labels as in Gordon parcellation. Group-level modularity partition was obtained using graph-theoretical consensus community detection. Gray peripheral bars around the connectograms indicate the number of connections per node. Flow diagram in the middle illustrates the reconfiguration of brain network modules from resting state (left) to the listening task (right). Modules are shown in vertical boxes with their heights corresponding to their connection densities. The streamlines illustrate how nodes belonging to a given module during resting state change their module membership under the listening task. See Figure 2-1 for the network analysis pipeline and Figure 2-2 for the activation maps.

Our network analysis revealed a significant increase in network modularity during the task compared to rest (permutation test; Cohen’s d = 0.3, p = 0.02; Fig. 2B, left). Notably, there was no difference in overall mean functional connectivity between resting-state and listening task networks (Cohen’s d = -0.01, p = 0.93). This underscores the importance of modular reconfiguration (i.e., changes in network topology) in adapting brain networks to the current task context, as opposed to mere alterations in overall functional connectivity. This finding aligns well with our previous study, where whole-brain network modularity increased during a dichotic listening task with concurrent speech (Alavash et al., 2019).

In an additional analysis, we explored the correlation between resting-state and task brain connectivity and network modularity separately. Reassuringly, both measures exhibited positive correlations across participants (connectivity: Spearman’s rho= 0.5, p < 0.01; modularity: rho= 0.61, p < 0.01), supporting the notion that both state- and trait-like determinants contribute to individuals’ brain network configurations (Geerligs et al., 2015). To Lending generality to all further results, we found no significant differences in network modularity between attending left versus attending right (permutation test; d = 0.04, p = 0.5), left versus front (d = 0.03, p = 0.75), or right versus front (d = 0.01, p = 0.86).

Subsequently, we explored whether the reconfiguration of whole-brain networks from rest to task could explain inter-individual variability in listening behavior (Fig. 1C). Using the same linear mixed effects model that demonstrated the beneficial effects of pitch difference and congruency of tone pairs on objective and subjective measures of behavior, we examined the direct and interactive effects of brain network modularity during the task in predicting listeners’ behavior. By incorporating single-trial accuracy as a binary predictor in our models, we were also able to assess test the interaction between network modularity and accuracy in predicting response speed or confidence. These models also included whole-brain mean connectivity as a between-subject covariate.

We found a significant main effect of network modularity in predicting single-trial response confidence (OR= 0.85, p < 0.01, log[Bayes factor(BF)] = 3.77; Fig. 3B). This result indicated that those listeners who showed higher brain network modularity during the task performed less confidently. However, prediction of confidence from network modularity did not differ between correct and incorrect judgments (modularity x accuracy interaction: OR= 1.09, p = 0.15). Taken together, these results provided initial evidence that, at the whole-brain level, reconfiguration of network modules is task-relevant, as it has the potential to explain individual differences in listeners’ response confidence.

The effects of pitch difference, congruency of the tone pairs, or the role of the lateral stream (i.e., to-be-attended or to-be-ignored) on single-trial confidence ratings did not moderate the whole­ brain network modularity effects. Adding an interaction term between these experimental conditions and brain network modularity did not significantly improve model fit (likelihood ratio tests; pitch difference: X^2^ = 0.54, p = 0.45; congruency: x^2^ = 0.01, p = 0.92; role of the lateral stream: x^2^ = 3.2, p = 0.07). Also, the interaction between rest and task modularity did not contribute to predicting single-trial accuracy or response speed (accuracy: OR= 1.02, p = 0.79; speed: = -0.02, p = 0.35).

In the following sections, we investigated the cortical origin of the modular reconfiguration outlined above: first, at the whole-brain level and second, at subnetwork and nodal levels in more detail. In the final section of the results, we will illustrate how these network dynamics can explain metacognitive differences in listening behavior.

### Reconfiguration of auditory and attentional-control networks during selective pitch discrimination

Fig. 3C provides a comprehensive overview of group-level brain networks, functional connectivity maps, and the corresponding connectograms (circular diagrams) during resting state (Fig. 3C, Left) and the listening task (Fig. 3C, Right). In Fig. 3, cortical regions defined based on Gordon parcellation (Gordon et al., 2016) are grouped and color-coded according to their module membership determined using the graph-theoretical consensus community detection algorithm (see *Materials and Methods: Network modularity).* When applied to the resting-state data in our sample (N = 40), the algorithm decomposed the group-averaged whole-brain network into six modules (Fig. 3C, Left). The network module with the highest number of nodes largely overlapped with the known default mode network (Fig. 3C, dark blue). Guided by our previous study (Alavash et al., 2019), our investigation was particularly focused on the module comprising mostly auditory nodes and a number of cinguloopercular nodes (Fig. 3C, yellow). For simplicity, similar to Alavash et al. (2019) we will refer to this module as the auditory module. Additional network modules included visual (Fig. 3C, cyan), somatomotor (Fig. 3C, light blue), dorsal attention (Fig. 3C, green), and frontoparietal module (Fig. 3C, red).

Fig. 3C illustrates how the configuration of the whole-brain network changed from rest to task. This network flow diagram illustrates whether and how cortical nodes belonging to a given network module during resting state (Fig. 3C, Left) changed their module membership during the listening task (Fig. 3C, Right). The streamlines depict how the nodal composition of network modules changed, indicating a functional reconfiguration. According to the streamlines, the auditory and dorsal attention modules showed the most prominent reconfiguration (Fig. 3C, yellow and green modules, respectively), while the other modules underwent only moderate reconfiguration. This reconfiguration aligns well with the results found in our previous study, in which participants completed resting state followed by a selective listening task with concurrent speech.

Specifically, this reconfiguration can be summarized by a nodal “branching” of the auditory module during selective pitch discrimination (Fig. 3C, left gray box) accompanied by the formation of a module comprised of cinguloopercular, somatomotor, dorsal attention, and visual nodes. Given the pure auditory nature of the listening task here, the less prominent changes in visual nodes seem unlikely to be behaviorally relevant to task performance. To rule this out, for our brain-behavior models, we conducted control analyses in which the main predictors were derived from visual or somatomotor networks (see section *Control analyses).* Taken together, we observed a reconfiguration of the whole-brain network that is dominated by alterations across auditory, cinguloopercular, and dorsal attention nodes.

Accordingly, we next identified the cortical origin of these network dynamics based on the underlying functional parcellation (Gordon et al., 2016). The identified subnetwork entails auditory, cinguloopercular, and dorsal attention nodes. We will refer to this large-scale cortical subnetwork collectively as the auditory attentional-control network. This subnetwork encompasses 96 nodes (of 333 whole-brain nodes). According to Fig. 3C, this subnetwork is, in fact, a conjunction across all auditory, cinguloopercular, and dorsal attention nodes. For the purpose of a more transparent illustration, the cortical map of the auditory attentional-control network is visualized in Fig. 4A.

**Figure 4.**
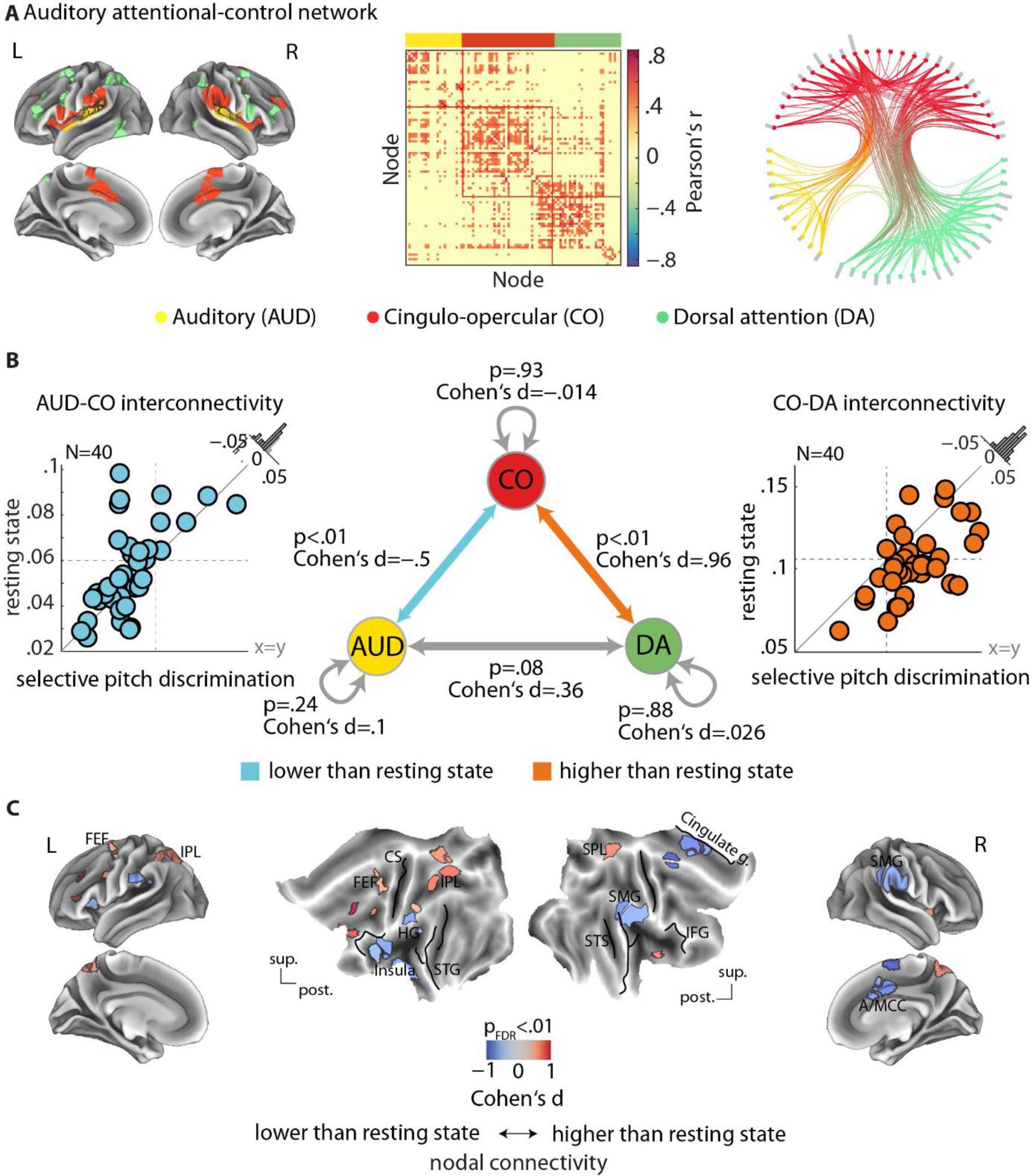
Modulation of connectivity across auditory and attentional-control networks. **(A)** The auditory attentional-control network composed of auditory (AUD), cinguloopercular (CO), and dorsal attention (DA) nodes. Nodes are identified according to their labels in Gordon parcellation. The correlation structure and connection pattern across these nodes are illustrated by group-level functional connectivity matrix and the connectogram (circular diagram), respectively. Gray peripheral bars around the connectogram indicate the number of connections per node. **(B)** Modulation of cinguloopercular connectivity with auditory and dorsal attention networks during the listening task. From rest to task, AUD-CO mean interconnectivity decreased (left plot), whereas CO­ DA interconnectivity increased (right plot). Mean connectivity across nodes within each network or between auditory and dorsal attention networks did not change significantly (middle graph, gray arrows). Histograms show the distribution of the change (task minus rest) across all 40 participants. (C) Modulation of nodal connectivity across the auditory attentional-control network. Nodal connectivity (also known as strength) was quantified as the sum of correlation values per node. The result was compared between task and rest per node using paired permutation tests and corrected for multiple comparison across nodes (FDR correction at significance threshold 0.01). Nodes exhibiting significant decrease in their connectivity during the listening task overlapped with the bilateral supramarginal gyri (SMG), the left anterior insula, and the anterior/middle cingulate cortices (A/MCC). Nodes showing significant increase in their connectivity overlapped with the superior parietal/intraparietal lobule (SPL/IPL) and the frontal eye fields (FEF). CS: central sulcus. HG: Heschl gyrus. /FG: inferior frontal gyrus. *STG/5:* superior temporal gyrus/sulcus.

### Modulation of interconnectivity across the auditory attentional-control network

To uncover the network dynamics driving the modular reconfiguration of the auditory attentional­ control network in adaptation to the listening task, we explored whether and how connectivity within and between auditory, cinguloopercular, and dorsal attention nodes changed during the task compared to rest.

The results are visualized in Fig. 4B. We found two distinct patterns of connectivity changes across the auditory attentional-control network. First, from rest to task, mean connectivity between auditory and cinguloopercular network (AUD-CO) was decreased significantly (Cohen’s d = -0.5, p < 0.01; Fig. 4B, Left). Second, mean connectivity between cinguloopercular and dorsal attention network (CO-DA) increased significantly (Cohen’s d = 0.96, p < 0.01; Fig. 4B, Right). Additionally, connectivity between auditory and dorsal attention network was increased, but the degree of this modulation did not reach the significance level (Cohen’s d = 0.36, p = 0.08; Fig. 4B, Bottom).

It is worth noting that mean connectivity within the auditory, cinguloopercular, or dorsal attention networks did not significantly differ from rest (Fig. 4B, gray self-connections). This observation underscores the importance of functional interplay between auditory and attentional­ control networks (i.e., interconnectivity) in regulating or contributing to the metacognitive aspects of listening behavior.

Interestingly, the interconnectivity between AUD-CO and CO-DA was not correlated across participants during both rest and task (rest: rho= -.14, p = 0.4, task: rho= -.03, p = 0.86). This suggests the presence of two dissociable large-scale networked systems shaping metacognitive performance during listening.

Subsequently, we performed a nodal analysis to identify the cortical regions responsible for the change of interconnectivity across the auditory attentional-control network. To this end, we used a simple graph-theoretical measure-nodal connectivity (or nodal strength)-which is defined as the sum of each node’s connection weights. Through permutation tests comparing nodal connectivity between rest and task, we found that connectivity of mainly supramarginal gyri, anterior insular, and anterior/mid-cingulate cortices was significantly decreased during the selective pitch discrimination task relative to rest (-1 < Cohen’s d < -0.53, p < 0.01, false discovery rate (FDR)-corrected for multiple comparisons across nodes; Fig. 4C, blue). Additionally, we found significant increase in connectivity in cortical nodes overlapping with parietal lobules and frontal eye-fields (0.43 < Cohen’s d < 1, p < 0.01; Fig.4C, red).

### Interconnectivity of auditory attentional-control network predicts metacognitive differences in listening behavior

As depicted in Fig. 4B, listeners exhibited clear inter-individual variability in the degree and direction of change in AUD-CO and CO-DA interconnectivity. Thus, we next explored whether these variabilities could explain inter-individual differences in listening behavior reported earlier (Fig. 1C). To this end, we examined the direct and interactive effects of individual mean interconnectivity during rest and task on single-trial response accuracy, speed, or confidence. Specifically, joint analysis of single-trial response accuracy with speed and confidence allowed us to investigate whether and how the interconnectivity dynamics underlie individual differences in metacognitive performance, namely response speed or confidence after correct/incorrect judgments. These analyses are based on the same linear mixed-effects models that revealed the beneficial effects of pitch difference and congruency on behavior as well as higher confidence and response speed on correct trials compared to incorrect trials ones.

The relationships between brain connectivity and behavior were examined separately for each AUD-CO and CO-DA network pair. In each model, brain regressors were included as between-subject covariates. In our models, we controlled for the activity level of each auditory, cinguloopercular, and dorsal attention network by incorporating mean beta estimates derived from nodal activation analysis (i.e., univariate general linear models) as additional between-subject covariates (see section *Control analyses)*.

Firstly, AUD-CO interconnectivity and response accuracy interacted in predicting listeners’ response confidence (OR = 0.87, p < 0.05, log(BF) = 2.44; Fig SA): Across listeners, lower AUD-CO interconnectivity was associated with lower confidence in their own perceptual judgments (main effect of AUD-CO interconnectivity on response confidence; OR = 1.13, p < 0.001), with this relationship being more pronounced following incorrect judgments compared to correct ones (incorrect: OR = 1.25, correct: OR = 1.09). Additionally, lower AUD-CO interconnectivity overall predicted slower decision-making during the task (= 0.08, p < 0.001, log(BF) = 13.73), with no difference in the degree of this effect between correct and incorrect judgments (interaction term: = -0.02, p = 0.67).

Secondly, the dynamics of CO-DA interconnectivity predicted individual differences in response confidence (Fig. SB). Specifically, we found a significant interaction between CO-DA interconnectivity and response accuracy in predicting listeners’ confidence (OR = 1.14, p < 0.05, log(BF) = 1.53; Fig. SB): Across listeners, higher CO-DA interconnectivity was associated with less confident behavior (main effect of CO-DA interconnectivity on response confidence: OR= 0.88, p < 0.001). This negative relationship was also more pronounced after incorrect judgments compared to correct ones (incorrect: OR= 0.8, correct: OR= 0.96).

In additional analyses, we investigated the possibility that these correlations might be driven by aspects related to response speed, potentially explaining the salience of the links observed in incorrect trials associated with slower responses. We thus included single-trial speed as an additional covariate in models predicting confidence. As expected, during trials on which listeners were confident in their decisions, their responses were also faster (main effect of response speed on single-trial confidence: OR = 2, p < 0.01). Importantly, however, the interactions between interconnectivity of each network pair and accuracy in predicting confidence remained almost unchanged when accounting for speed in the models (AUD-CO model: OR= 0.87, p = 0.01; CO-DA: OR = 1.12, p = 0.07). In addition, in models predicting response speed, the interactions between inter-connectivity of each network pair and accuracy were not significant (AUD-CO: = -0.02, p = 0.67; CO-DA: = 0.02, p = 0.63). Taken together, the salience of the links observed in incorrect trials was specific to prediction of confidence, and not speed.

After uncovering the link between the interconnectivity dynamics of the auditory attentional­ control network and individual listening behavior, we proceeded to identify which cortical regions within this network could explain metacognitive differences in listening behavior. For this purpose, we estimated a linear mixed-effects model for each node within the auditory attentional-control network, similar to the brain-behavior analyses reported above. In these models, however, we substituted the predictor representing the mean interconnectivity of the network pairs (i.e., AUD-CO or CO-DA mean interconnectivity) with nodal connectivity of each region within the auditory attentional-control network. Consequently, the p-values obtained for the interaction between nodal connectivity and response accuracy were adjusted for multiple comparisons across nodes (FDR correction at a significance threshold 0.01).

We found significant interactions between response accuracy and nodal connectivity of brain regions overlapping with the auditory/insular, superior parietal, and middle cingulate cortices (Fig. 6A). The direction of these interactions differed between two sets of nodes. Specifically, cortical nodes near the HeschI gyrus (HG), anterior insula, and middle cingulate cortex (MCC) exhibited odds ratios smaller than one, indicating a relation between their nodal connectivity and listeners’ response confidence consistent with the effect found based on mean AUD-CO interconnectivity (Fig. 6A, blue regions). In contrast, cortical nodes near the bilateral superior parietal lobule (SPL) and left inferior parietal lobule (IPL) showed odds ratios larger than one, indicating a relationship between their nodal connectivity and listeners’ response confidence consistent with the effect found based on mean CO-DA interconnectivity (Fig. 6A, red regions).

To illustrate, interactions found in the left IPL and right MCC are shown in Fig 6B. Consistent with the results obtained based on mean AUD-CO interconnectivity (Fig. SA), we found a significant interaction between nodal connectivity of the right MCC and response accuracy in predicting listeners’ confidence (OR = 0.76, p < 0.001, log(BF) = 16.4; Fig 6B, left). This finding indicates that lower nodal connectivity of the right MCC was associated with less confident responses across listeners, but this relationship was specific to incorrect judgments (incorrect: OR= 1.2; correct: OR= 0.97). Additionally, in line with the results obtained based on mean CO-DA interconnectivity (Fig. SB), we found a significant interaction between nodal connectivity of the left IPL and response accuracy in predicting listeners’ confidence (OR= 1.3S, p < 0.001, log(BF) = 17.1; Fig 6B, right): Higher nodal connectivity of IPL was associated with less confident responses across listeners, but this relationship was specific to incorrect judgments compared to correct ones (incorrect: OR = 0.77, correct: OR = 0.98).

### Control analyses

Our findings illustrate that the dynamics of interconnectivity across the auditory attentional-control network could partly explain individual differences in subjective listening behavior. These dynamics were expressed as modulation of mean connectivity between network pairs defined using average Pearson correlations, a commonly used measure of functional connectivity. Accordingly, one question is, to what extent the brain-behavior relations observed here are driven by changes in cortical activity level rather than changes in correlation strength. In addition, from rest to task, the somatomotor and visual networks also underwent a moderate reconfiguration (Fig. 3C). Thus, another question is whether interconnectivity with somatomotor or visual network can predict listeners’ behavior. Therefore, we conducted a set of control analyses to ensure that the brain­ behavior findings are not trivial.

First, all of the brain-behavior models have been adjusted for individual mean cortical activity of each auditory, cinguloopercular, and dorsal attention network by including between-subject covariates of mean beta estimates derived from activation analysis (Fig. 7). Also, none of these potential confounding predictors on behavior proved to be significant (Table 1A).

**Table 1.** Summary tables of control analyses.

Prediction of response confidence from mean activity level of cortical networks.
| Network | Event onset | Main effect of mean beta estimate |
| --- | --- | --- |
| AUD | Spatial cue | OR = 0.6<br>p = 0.53 |
| CO | Spatial cue | OR = 0.8<br>p = 0.7 |
| DA | Spatial cue | OR = 1.9<br>p = 0.39 |
| AUD | Tone sequence | OR = 1.3<br>p = 0.7 |
| CO | Tone sequence | OR = 1.6<br>p = 0.76 |
| DA | Tone sequence | OR = 0.75<br>p = 0.77 |

**B.** Prediction of response confidence from mean interconnectivity of other networks.
| Network pair | Accuracy × Interconnectivity |
| --- | --- |
| SM-AUD | OR = 0.86<br>p = 0.003 |
| SM-CO | OR = 0.97<br>p = 0.63 |
| SM-DA | OR = 1.08<br>p = 0.24 |
| VIS-AUD | OR = 0.89<br>p = 0.1 |
| VIS-CO | OR = 0.96<br>p = 0.72 |
| VIS-DA | OR = 0.93<br>p = 0.53 |

AUD: auditory; CO: cinguloopercular; DA: dorsal attention; SM: somatomotor; VIS: visual

Second, we tested additional brain-behavior models in which the connectivity regressors were defined based on interconnectivity between each somatomotor or visual network with the auditory, cinguloopercular, or dorsal attention network. Identical to our main analysis, these models were tested separately for each network pair in predicting single-trial response confidence. The results are provided in Table 1B. We only found a significant interaction between response accuracy and interconnectivity of somatomotor and auditory networks in predicting listeners’ response confidence (OR= 0.86, p < 0.01). This result indicates that lower AUD-SM interconnectivity predicted lower response confidence on incorrect trials, similar to the results obtained based on AUD-CO interconnectivity.

In additional analyses we investigated the degree to which individual metacognitive differences might be related to hearing sensitivity, pitch discrimination ability, or task difficulty. To this end, we included two additional covariates in our models: (1) individuals’ hearing thresholds measured using PTA (averaged between left and right ear within frequencies 0.5, 1, 2, and 4 kHz), and (2) listeners’ ratings of their own musical ability (ranging from 1 for expert to 6 for naYve; M±SD: 3±1.7) which we collected at the end of the experiment. Neither of these measures had a significant effect on response confidence (PTA: OR = 1.25, p = 0.78; musical experience: OR = 0.5, p = 0.08). In addition, across listeners, neither of the correlations between overall mean response accuracy or pitch difference and mean response confidence were significant (accuracy: Spearman’s rho = -0.12, p = 0.48; pitch difference: rho = 0.1, p = 0.54). These suggest that, across listeners, metacognitive differences were unlikely to be related to differences in task difficulty. Importantly, our main findings, .e., the interaction between brain inter-connectivity and accuracy in predicting confidence (Fig. 5) remained significant after these covariates were accounted for.

**Figure 5.**
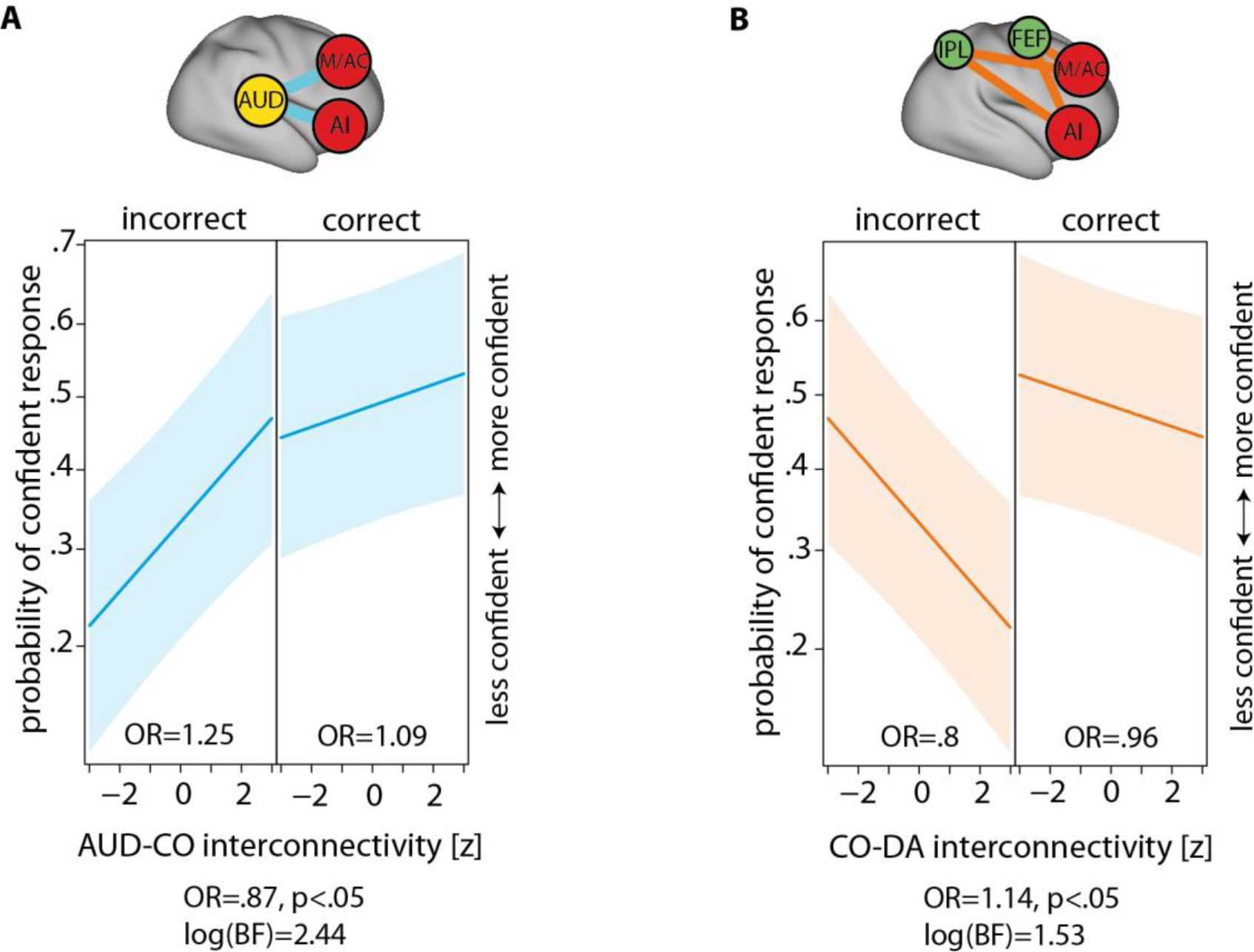
Prediction of individual listening confidence from interconnectivity dynamics of auditory attentional-control network. **(A)** Prediction of single-trial response confidence from mean AUD-CO interconnectivity during the selective pitch discrimination task. Across listeners, lower AUD-CO interconnectivity predicted less confident responses during the task, the degree of which was more pronounced during incorrect judgments compared to correct ones. **(B)** Prediction of single-trial response confidence from mean CO-DA interconnectivity. Across listeners, higher CO­ DA interconnectivity predicted less confident responses during the task, the degree of which was stronger during incorrect judgments compared to correct ones. Experimental variables including single-trial pitch differences, congruency of tone pairs, and location of spatial cue have been accounted for in the models. Shaded areas show two-sided parametric 95% Cl. OR: odds ratio parameter estimates from generalized linear mixed-effects model. A/: Anterior insula. *FEF:* Frontal eye-fields. */PL:* Inferior parietal lobule. *M/AC:* Mid-/anterior cingulate. See Table 4-1 for the summary of control analysis.

## Discussion

The aim of our study was to elucidate the variability among individuals in the subjective aspects of listening behavior by examining the dynamics of cortical networks related to audition and cognitive control. Importantly, we rigorously controlled and evaluated listening behavior objectively using response accuracy or speed, while also allowing listeners to subjectively reflect on their own perceptual decisions by reporting their response confidence. Our focus was on the considerable variability observed in this metacognitive aspect of listening behavior.

First, by investigating the modular reconfiguration of brain networks from a resting state to task engagement, we identified the auditory attentional-control network. Subsequently, across this network and on an individual level, we found that task performance influenced the interconnectivity between the cinguloopercular network and each of the auditory and dorsal attention networks. The results obtained from our brain-behavior analyses supported the functional significance of these interconnectivity dynamics.

In summary, our findings suggest that decreased interconnectivity between auditory and cognitive control network (AUD-CO) and increased interconnectivity between cognitive control and dorsal attention networks (CO-DA) are indicative of heightened metacognitive sensitivity, specifically lower confidence in responses following incorrect judgments. These network configurations may facilitate listeners’ accurate self-assessment of their auditory perceptual decision.

Overall, our results indicate that in situations where listeners make metacognitive judgments about auditory information, the brain networks adaptively reconfigure towards a more introspective, control mode. This proposition will be further elaborated upon in subsequent sections.

### Metacognitive facets of adaptive listening behavior

Our listening paradigm exemplifies a situation where each listener performed well in terms of first­ order performance, yet there was variation in self-awareness regarding one’s own perception among listeners. This underscores an important aspect in the study of adaptive listening behavior, which can be formalized within the scope of metacognition as “metacognitive performance”, defined as the ability to introspect about the correctness of one’s own thoughts and actions (Fleming et al., 2012a). This additional dimension could complement other factors such as motivation or effort in the study of listening behavior (Pichora-Fuller et al. 2016; Peelle 2018). An intriguing question arises: how do metacognitive judgments relate to an individual’s motivation in challenging listening situations? The present experimental design does not allow measuring motivation, as it was not the main scope of the study (see, e.g., Kraus et al. 2024). An important distinction here, however, would be the introspective, thus self-monitoring nature of meta-listening.

While previous research has linked individual differences in brain structure and function to such variations (Ais et al., 2016), our findings specifically associate this variability with individual differences in the interconnectivity dynamics of two cortical networked systems, as discussed below.

### Interconnectivity of auditory and cinguloopercular networks regulates perceptual decision confidence during listening

Our first main finding was that lower connectivity between auditory and cinguloopercular network predicted slower and less confident responses during the listening task, the latter link being more pronounced for incorrect judgments (Fig. SA). This result is in line with the increase in network modularity (equivalently, functional segregation) observed on the whole-brain level, and in particular the prediction of less confident responses from greater functional segregation of brain networks (Fig. 3B).

Previous fMRI studies consistently demonstrate that cinguloopercular activity correlates with word recognition in speech-in-noise experiments (Vaden et al., 2013; Eckert et al., 2016; Vaden et al., 2019). Recently, using a perceptual decision-making model, it has been shown that cinguloopercular activity contributes to criteria adjustments to optimize speech recognition performance (Vaden et al., 2022). In parallel, visual studies have shown that cinguloopercular network displays sustained activity during perceptual processing (Sestieri et al., 2014), with its activity level relating to the speed of stimulus detection (Coste and Kleinschmidt, 2016). These findings can be interpreted in the light of the proposed role for cinguloopercular cortex in performance monitoring, and more generally, adaptive control (Dosenbach et al., 2008; Power and Petersen, 2013; Neta et al., 2014; Neta et al., 2017).

Interestingly, previous work on neural correlates of metacognition during decision-making behavior support the role of frontal brain areas in subjective sense of decision confidence (Vaccaro and Fleming, 2018). Specifically, studies have found that explicit confidence estimates are tracked in cortical areas including medial/rostrolateral prefrontal cortex (De Martino et al., 2013; Hebart et al., 2016), and dorsal anterior cingulate cortex (Fleming et al., 2012b; Fleming et al., 2014). Additionally, the medial prefrontal cortex has been linked to inter- and intra-individual variation in explicit confidence estimates (Bang and Fleming, 2018). Direct evidence comes from an animal study which associated the orbitofrontal cortex with confidence-based behaviors (Kepecs et al., 2008). In our data, this is supported by a link between connectivity of a node within the right middle cingulate cortex and listeners’ response confidence on incorrect trials (Fig. 6B).

**Figure 6.**
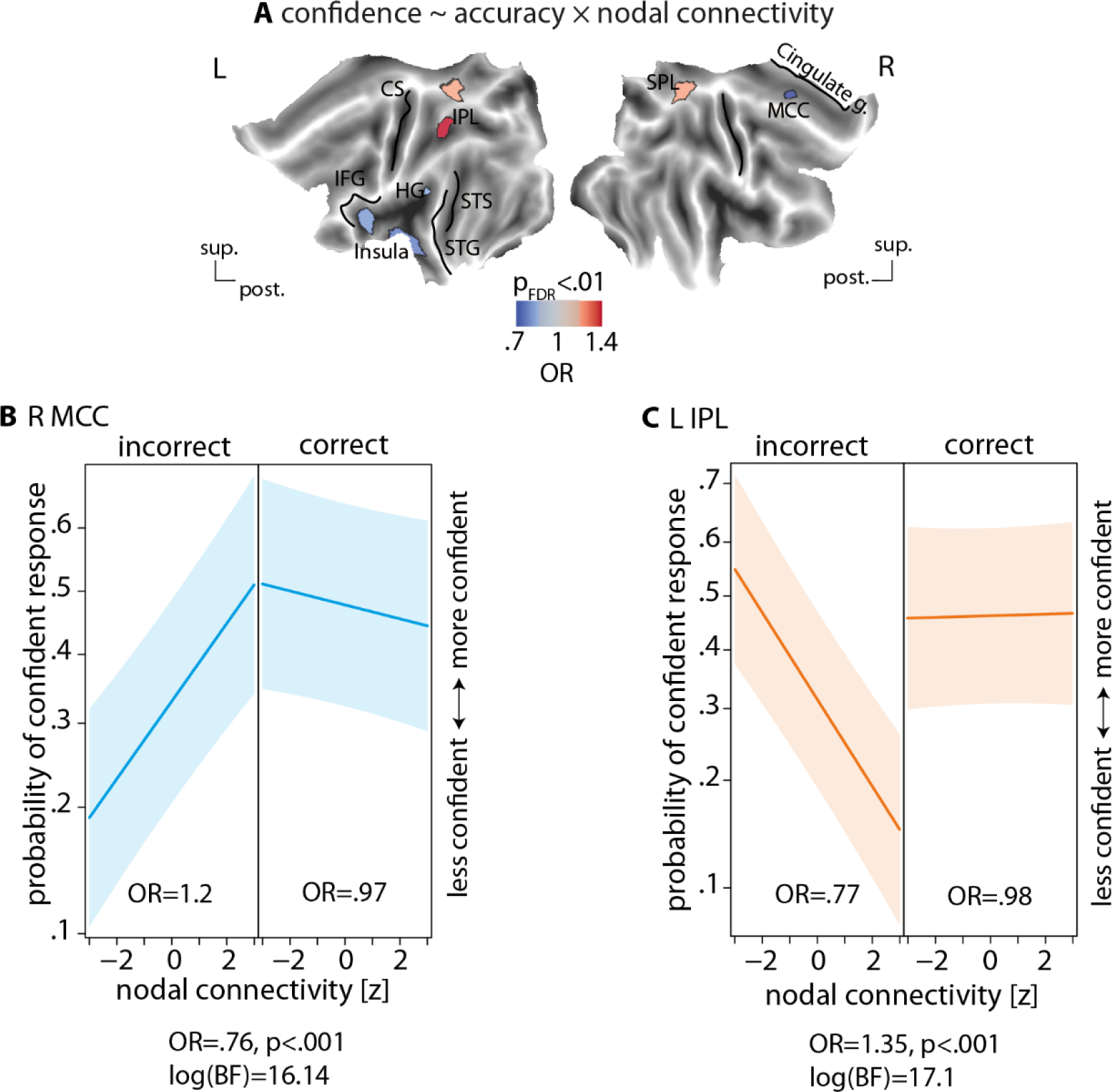
Prediction of individual listening confidence from nodal connectivity of the auditory attentional-control network. **(A)** Interaction between response accuracy and nodal connectivity in predicting single-trial confidence using generalized linear mixed effects models. Experimental variables including single-trial pitch differences, congruency of tone pairs, and location of spatial cue have been accounted for in the models. The models were tested for each node of the auditory attentional-control network. The p-values obtained for the interaction term were corrected for multiple comparisons across nodes (FDR correction at significance threshold 0.01).Two sets of nodes showed significant interactions: regions near the HeschI gyrus (HG), the anterior insular (Al), and the middle cingulate cortex (MCC) showed odds ratios smaller than one (indicating a more positive slope during incorrect than correct judgments), and regions overlapping with the superior/inferior parietal lobules (SPL/IPL) showed odds ratios larger than one (indicating a more negative slope during incorrect than correct judgments). **(B)** To illustrate, resolving the interaction found at the right MCC indicated that lower connectivity of this node predicted lower confidence following incorrect judgments. (C) Resolving the interaction found at the left IPL indicated that higher connectivity of this node predicted lower confidence following incorrect trials. Shaded areas show two-sided parametric 95% Cl. OR: odds ratio parameter estimates from generalized linear mixed-effects model. CS: central sulcus. HG: HeschI gyrus. /FG: inferior frontal gyrus. *STG/5:* superior temporal gyrus/sulcus.

**Figure 7.**
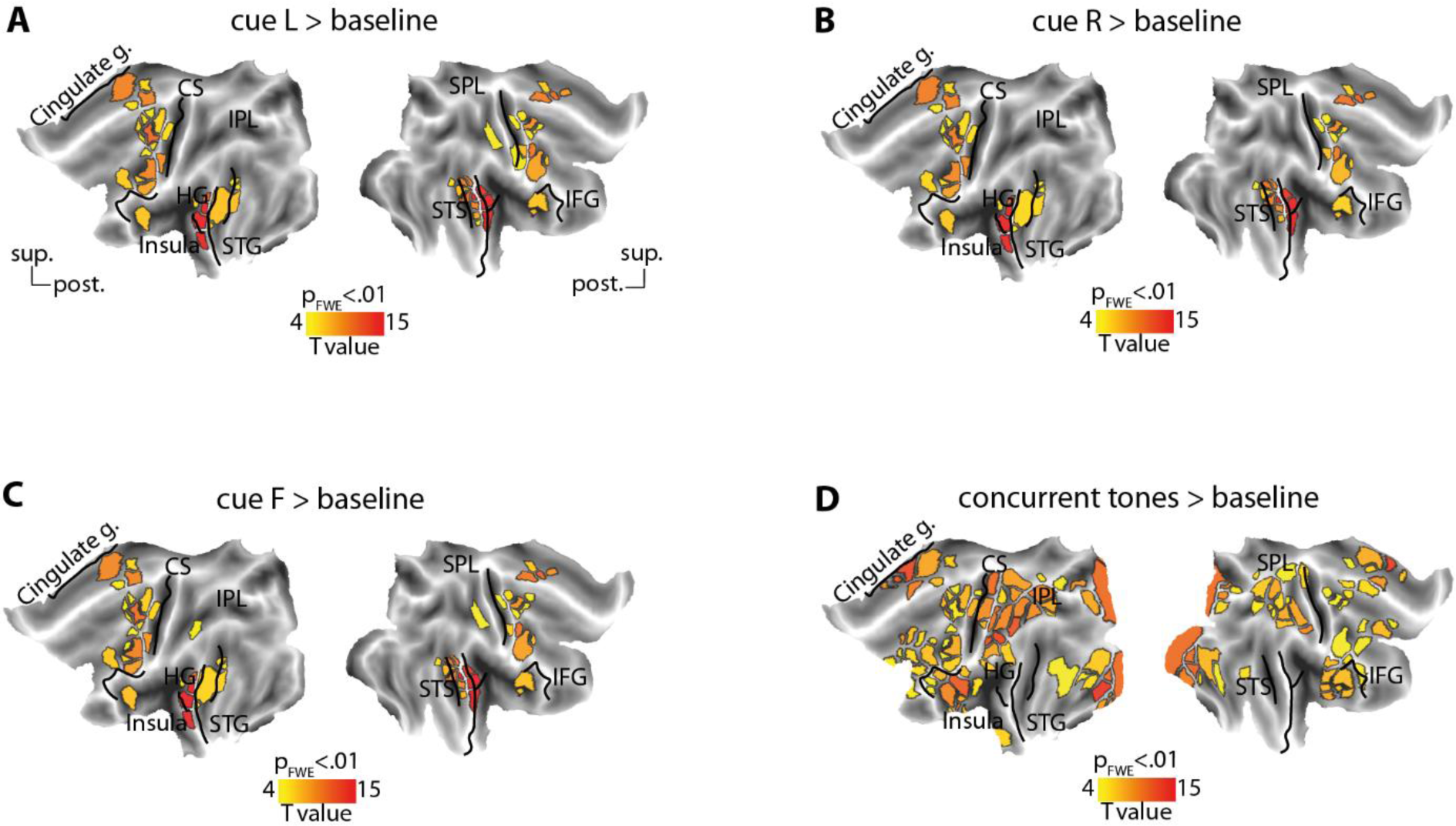
Group-level activation maps of task events. To derive individual estimates of cortical activity in response to different task events, an activation analysis using univariate general linear models (GLM) was conducted and individual statistical parametric maps were obtained. The GLM model was estimated per cortical node of the parcellation used for connectivity analysis. The design matrix included the following regressors: cue left, cue right, cue front, concurrent tones (modeled as 1-second epoch), and button press. The onset of these events in milliseconds were convolved with canonical hemodynamic response function (HRF) with temporal resolution of 1 millisecond, and the results were downsampled to one TR (i.e., 1s). These regressors were used to predict the same nodal timeseries used for connectivity analysis, i.e., the mean BOLD timeseries recovered following nuisance regression and band-pass filtering per node. Finally, group-average statistical maps were obtained by submitting the beta estimates to one-sided t-tests (against implicit baseline) per node, and the results were thresholded at p<0.01 following correction for multiple comparisons across nodes using family-wise error correction (FEW) procedure. The resulting corrected maps are visualized for different conditions in (A) cue left, (B) cue right, (C) cue front, and (D) concurrent tone presentation. None of the differential contrasts across the cue conditions, i.e., left vs. right or left/right vs. front, survived the significance threshold.

The functional overlap between the cinguloopercular network, adaptive control, and frontal cortical areas representing decision confidence aligns with the notion of a domain-general role of the cinguloopercular network in the neural representation of metacognitive performance (Baird et al., 2013; Rouault et al., 2018; Rouault et al., 2023).

### Interconnectivity of cinguloopercular and dorsal attention networks supports metacognition during selective listening

As second main finding, higher interconnectivity between cinguloopercular and dorsal attention networks predicted listeners’ lower confidence during the task after incorrect judgments (Fig. SB). Selective attention, especially in visual and spatial domains, is known to recruit dorsal attention network (Corbetta et al., 2008; Petersen and Posner, 2012; Posner, 2012; Vossel et al., 2014). Neuroimaging studies have shown that spatial auditory attention recruits some of the same cortical regions as visuospatial attention (Lee et al., 2014). In addition, it has been shown that cinguloopercular network is flexibly linked with dorsal attention network during visuospatial or memory search, respectively, consistent with the characterization of the cinguloopercular network as a domain-general task-control system (Sestieri et al., 2014). Studies in non-human animals have also associated the canonical nodes of dorsal attention network, namely inferior parietal sulcus and frontal eye fields, to metacognitive performance during visual-spatial tasks (Kiani and Shadlen, 2009; Middlebrooks and Sommer, 2012).

### Comparison to previous work

Our study stands out in two significant ways. Firstly, unlike previous studies which focused on word recognition task or categorical perceptual decision-making tasks, our task required listeners to flexibly shift and maintain auditory attention on a trial-by-trial basis, while monitoring their perception and reporting their decision confidence. Thus, our task design allowed a combined analysis of both objective and subjective behavior during adaptive listening. To our knowledge, this is the first study that casts the problem of adaptive listening behavior into the flourishing framework of metacognition in perceptual decision-making.

Secondly, while previous studies on metacognition often establish correlations between brain activity and a latent state of uncertainty estimated using computational models (Walker et al., 2023), our study instead took a connectionist approach and investigated individual differences in meta­ listening within auditory and higher-order attentional-control networks.

### Limitations and future directions

The interconnectivity dynamics uncovered in our study suggest a potential role of network hubs in guiding individuals’ meta-listening (cf. Gratton et al., 2017). Control networks do not function in isolation but rather exhibit context-dependent dynamic interactions during adaptive behaviors (Menon and D’Esposito, 2022).

On a neurophysiological level, previous research from our lab has provided important insights into how the dynamics of low-frequency neural oscillations underlie individual differences in objective measures of listening behavior (Alavash et al., 2017; Wöistmann et al., 2019; Alavash et al., 2021; Tune et al., 2021). Recent studies on the neural correlates of metacognition during perceptual decision-making suggest a link between pre-stimulus neural activity in the alpha frequency range and confidence (Kayser et al., 2016; Wöstmann et al., 2019). Given the associations between modulations in alpha oscillations and the activity of cinguloopercular and dorsal attention networks (Sadaghiani and Kleinschmidt, 2016), it is conceivable that alpha oscillations organize the interconnectivity dynamics found in our study.

Lastly, our findings hold potential clinical implications. Age-related hearing loss often affects the neural and behavioral implementation of auditory selective attention to varying degrees among individuals (Dai et al., 2018; Tune et al., 2021). Objective measures of listening behavior, however, are increasingly recognized to not fully capture the problems that hearing impaired individuals experience in demanding listening situations (Wostmann et al., 2021). To understand, predict, or compensate this deficit at the individual level, it is essential to find neural signatures of also subjective listening performance. As this initial study demonstrates, how hearing loss might affect the capacity to monitor or calibrate one’s own listening behavior (i.e., meta-listening) is uncharted. Our study suggests interareal connectivity of brain networks as a first approximation for addressing this question at both the behavioral and neural levels (Waters et al., 2012).

## Acknowledgements

Research was supported by German Research Foundation (DFG) grant AL 2408/1-1 to M.A. The task paradigm had been developed jointly by author M.A. and Malte Wostmann. Anne Herrmann helped acquire the fMRI data. Proofreading the manuscript was assisted by ChatGPT 3.5.

## Data availability

The complete dataset associated with this work, including MRI data in BIDS format, is publicly available on the Open Science platform OSF at https://osf.io/a9cte/.

